# *Leishmania infantum* xenodiagnosis from vertically infected dogs reveals significant skin tropism

**DOI:** 10.1101/2021.04.08.438940

**Authors:** Breanna M. Scorza, Kurayi Mahachi, Arin C. Cox, Angela J. Toepp, Adam Leal-Lima, Anurag Kumar Kushwaha, Patrick Kelly, Claudio Meneses, Geneva Wilson, Katherine N. Gibson-Corley, Lyric Bartholomay, Shaden Kamhawi, Christine A. Petersen

## Abstract

**Background:** Dogs are the primary reservoir for human visceral leishmaniasis due to *Leishmania infantum*. Phlebotomine sand flies maintain zoonotic transmission of parasites between dogs and humans. A subset of dogs is infected transplacentally during gestation, but at what stage of the clinical spectrum vertically infected dogs contribute to the infected sand fly pool is unknown.

**Methodology/Principal Findings:** We examined infectiousness of dogs vertically infected with *L. infantum* from multiple clinical states to the vector *Lutzomyia longipalpis* using xenodiagnosis and found that vertically infected dogs were infectious to sand flies at differing rates. Dogs with moderate disease showed significantly higher transmission to the vector than dogs with very mild or severe disease. We documented a substantial parasite burden in the skin of vertically infected dogs by RT-qPCR, despite these dogs not having received intradermal parasites via sand flies. There was a highly significant correlation between skin parasite burden at the feeding site and sand fly parasite uptake. This suggests dogs with high skin parasite burden contribute the most to the infected sand fly pool. Although skin parasite load and parasitemia correlated with one another, the average parasite number detected in skin was significantly higher compared to blood in matched subjects. Thus, dermal resident parasites were infectious to sand flies from dogs without detectable parasitemia.

**Conclusions/Significance:** Together, our data implicate skin parasite burden and earlier clinical status as stronger indicators of outward transmission potential than blood parasite burden. Our studies of a population of dogs without vector transmission highlights the need to consider canine vertical transmission in surveillance and prevention strategies.

**AUTHOR SUMMARY:** Sand flies transmit *Leishmania* parasites between infected dogs and humans leading to the life-threatening tropical disease Visceral Leishmaniasis (VL). Identifying which dogs transmit parasites well to sand flies is important to curb disease spread. The offspring of both dogs and humans can also be infected vertically while *in utero*. Despite this, the infectiousness of dogs that receive parasites *in utero* to sand flies has not been thoroughly investigated. Thus, we allowed sand flies to feed on a group of vertically infected dogs at varying stages of VL disease severity and measured sand fly parasite uptake. We found vertically infected dogs were readily able to transmit parasites to the sand flies. Dogs that were most infectious had moderate clinical disease and relatively high levels of parasite infection in their blood and skin. However, the level of skin infection was significantly higher than that observed in the blood, and the skin parasite load had the strongest correlation with sand fly parasite uptake. This implicates the skin may be an underappreciated driver of canine infectiousness to the sand fly vector. In addition, this work highlights that vertically infected dogs are very important parts of the transmission cycle and must be considered in all public health efforts addressing VL.

## INTRODUCTION

Visceral leishmaniasis (VL) is caused by the protozoan *Leishmania infantum*, killing an estimated 30,000 people annually [1]. In *L. infantum* endemic areas, dogs are the primary domestic reservoir driving zoonotic VL [2-4]. Vector transmission by phlebotomine sand flies is the predominant mode of parasite transmission; however, transplacental transmission has been demonstrated in both humans and dogs with VL [4]. In the United States, *L. infantum* is enzootic in hunting dogs and vertical transmission alone maintains infection within this population [21, 22]. As human VL is inextricably linked to canine infection, understanding factors underlying transmission from vertically *L. infantum*-infected dogs to sand flies is of utmost public health concern.

Epidemiological modeling and experimental findings support that a fraction of *L. infantum* infected dogs, termed ‘super-spreaders’, disproportionately contribute to the pool of infectious sand flies [5, 6]. Several groups have applied xenodiagnosis, the gold standard for measuring pathogen transmissibility from an infected host to an insect vector, in an effort to identify risk factors associated with increased canine transmission to sand flies [7-9]. Parasitemia and clinical status are the most commonly identified correlates of infectiousness to sand flies [10-12]. However, widely variable amounts of parasite uptake after xenodiagnosis between sand flies fed on the same host indicates other microenvironmental factors influence this interaction [13].

Importantly, sand flies are telmophages, feeding from blood pools formed by lacerating mammalian skin with a serrated proboscis [13]. This type of feeding allows parasite uptake from dermal blood vessels, as well as disrupted dermal-resident parasitized cells. Recent findings using xenodiagnosis have implicated canine skin parasite load as a critical correlate of canine infectiousness to sand flies [5, 12, 14, 15].

When sand flies egest *L. infantum* into the skin of naïve canine hosts, parasite depots persist at the skin inoculation site [16]. This finding could indicate skin resident parasites are remnants of previous vector bite-site infections. In contrast, congenital transmission of *L. infantum* occurs via a hematological route. This raises the question of whether vertically infected dogs attain a skin parasite burden high enough to influence transmission in the absence of receiving intradermal parasite inoculations from sand flies. Dermotropism of *L. infantum* in dogs known to be infected vertically has not been investigated. Specifically, it is not known whether the same mechanisms that govern parasite transmission during vector transmission will also dictate parasite localization in vertically infected dogs and their ability to transmit out to the sand fly vector.

To address these questions, we assayed skin parasite burden and performed xenodiagnosis on a cohort of dogs vertically infected with *L. infantum* at various stages of clinical disease. We found significant dermotropism of *L. infantum*, highly correlated with transmission to sand flies. This confirms skin parasite infection is linked to parasite transmission from both congenitally and vector infected dogs.

## MATERIALS AND METHODS

### Animal Cohort

Canine subjects in this study were donated to the University of Iowa after signed informed consent was obtained. *L. infantum* infection of all dogs occurred naturally *in utero*. All animal use involved in this work was performed under the supervision of licensed and where appropriate board-certified veterinarians according to International AAALAC accreditation standards and institutional IACUC approval. The Leishvet clinical scores and demographics of the 16 dogs used in this study are provided (Supplemental Table 1.)

### Leishmaniosis clinical classification

On the day of xenodiagnosis, a veterinarian conducted a physical exam on each dog for clinical signs of leishmaniosis; whole blood and serum were collected. Whole blood was subjected to complete blood count (IDEXX Laboratories Inc.) and DNA isolation using the QIAamp Blood DNA Mini Kit according to manufacturer instructions (Qiagen). Real Time-quantitative PCR (RT-qPCR) for *Leishmania* ribosomal DNA was performed as previously described [17]. Serum was used to perform Dual-Path Platform ® Canine Visceral Leishmaniosis (DPP) serological analysis (ChemBios) detecting antibodies against recombinant *L. infantum* rK28 antigen. ChemBios Micro Reader was used to determine serological status with a reader value >10 considered as seropositive [18]. Serum chemistry panels were performed (IDEXX Laboratories Inc.), Supplemental Table 2. In some cases, urine was analyzed by refractometer. The results of physical exam and these tests were combined to clinically stage leishmaniosis as: subclinical (stage 1), mild (stage 2), moderate (stage 3), or severe disease (stage 4) according to the LeishVet Guidelines [19].

### Sand Flies and Xenodiagnosis

All sand fly handling was performed using the BEI Resources WRAIR Care and Maintenance of Phlebotomine Sand Flies guidelines. Colony-bred *Lutzomyia longipalpis* sand flies were incubated in a humidified chamber at 26**°**C with access to 30% sucrose solution. Sand flies were starved 12 hrs prior to xenodiagnosis. For feeding, ∼30-40 female sand flies were gently aspirated along with males at an approximate ratio of 3:1 and transferred into 2-inch diameter polycarbonate feeding cups covered on one side with a fine screen-mesh.

Dogs were sedated with dexmedetomidine (Zoetis Inc.). Heartrate and respiratory rate were monitored throughout procedure. Dogs were placed within an enclosed mesh chamber, and two feeding cups per dog were placed on the axillary region and inner pinna for 30 minutes.

Following feeding, engorged blood-fed female sand flies were separated and incubated for 48 hours at 26**°**C in a humidified chamber with access to sucrose solution. Individual blood-fed female flies were separated into Eppendorf tubes containing 100 uL of DNA lysis buffer (Gentra Puregene Tissue Kit, Qiagen) and stored at -20**°**C.

### Tissue isolation

Following xenodiagnosis and humane euthanasia per AVMA guidelines, a full necropsy was performed by a board-certified veterinary pathologist. Canine spleen samples were obtained, flash frozen on dry ice, and stored at -80**°**C until DNA isolation. Skin biopsies were taken from axillary and pinnal sand fly feeding sites as well as contralateral, unfed sites from each study animal using 6 mm punch biopsies (Integra) and placed into zinc formalin for histology or frozen in saline at -80**°**C for nucleic acid isolation.

### DNA isolation and Real Time-qPCR

For preparations of standard curves, low passage *Leishmania infantum* promastigotes (US/2016/MON1/FOXYMO4), originally isolated from a hunting dog with VL, were cultured in complete hemoflagellate-modified minimal essential medium with 10% heat-inactivated fetal calf serum at 26 °C and stationary phase parasites were enumerated by hemacytometer [20]. For sand fly parasite quantification, control female *Lu. longipalpis* sand flies fed on uninfected rabbit blood, incubated 48hrs, then were spiked with known quantities of *L. infantum* promastigotes before DNA isolation. For *Leishmania* quantification in host tissues, a standard curve was created by spiking known quantities of *L. infantum* promastigotes into uninfected canine blood and normalized by input DNA concentration.

Sand fly and canine sample DNA were isolated with the Gentra Puregene Tissue Kit (Qiagen). For *L. infantum* detection by RT-qPCR, *Leishmania* small-subunit rRNA-specific probe (5’-[6-FAM]-CGGTTCGGTGTGTGGCGCC-MGBNFQ-3’) and primers (forward, 5’-AAGTGCTTTCCCATCGCAACT-3’; reverse, 5’-GACGCACTAAACCCCTCCAA-3’) were used as previously described [17]. An exponential regression was constructed from the standard curve and used to convert Ct into parasite equivalents per sand fly or per 2.5 ug mammalian DNA.

### Statistical Analyses

Normality of data was assessed using the D’Agostino-Pearson test. For comparisons between tissues obtained from the same subject, Wilcoxon matched-pairs signed rank test was performed. For comparisons between subjects, Kruskal-Wallis ANOVA was performed. When appropriate, Dunn’s post-test was used for multiple comparisons. Spearman correlation was computed for correlation analyses. For all analyses, significance is: *p≤0.05; **p<0.01; ***p<0.001; ****p<0.0001.

## RESULTS

### Vertically infected dogs harbor significant skin parasite burden

In *L. infantum* endemic areas with vector predominant transmission, sand flies inoculate parasites into host skin, which then disseminate systemically but leave dermal foci of infection at the original bite site(s) [16]. We sought to investigate whether *L. infantum* parasites obtained *in utero* via the bloodstream which lack these original foci of parasites in the skin would also be dermotropic or remain concentrated in viscera. We analyzed parasite burdens in the blood, spleen, and skin in vertically infected dogs from this cohort at various clinical stages of VL (Figure 1). We utilized the LeishVet guidelines, which consider clinical physical signs and clinicopathologic laboratory findings, to stage canine VL clinical severity: subclinical (stage 1), mild (stage 2), moderate (stage 3), or severe disease (stage 4) [19].

**Figure 1.**
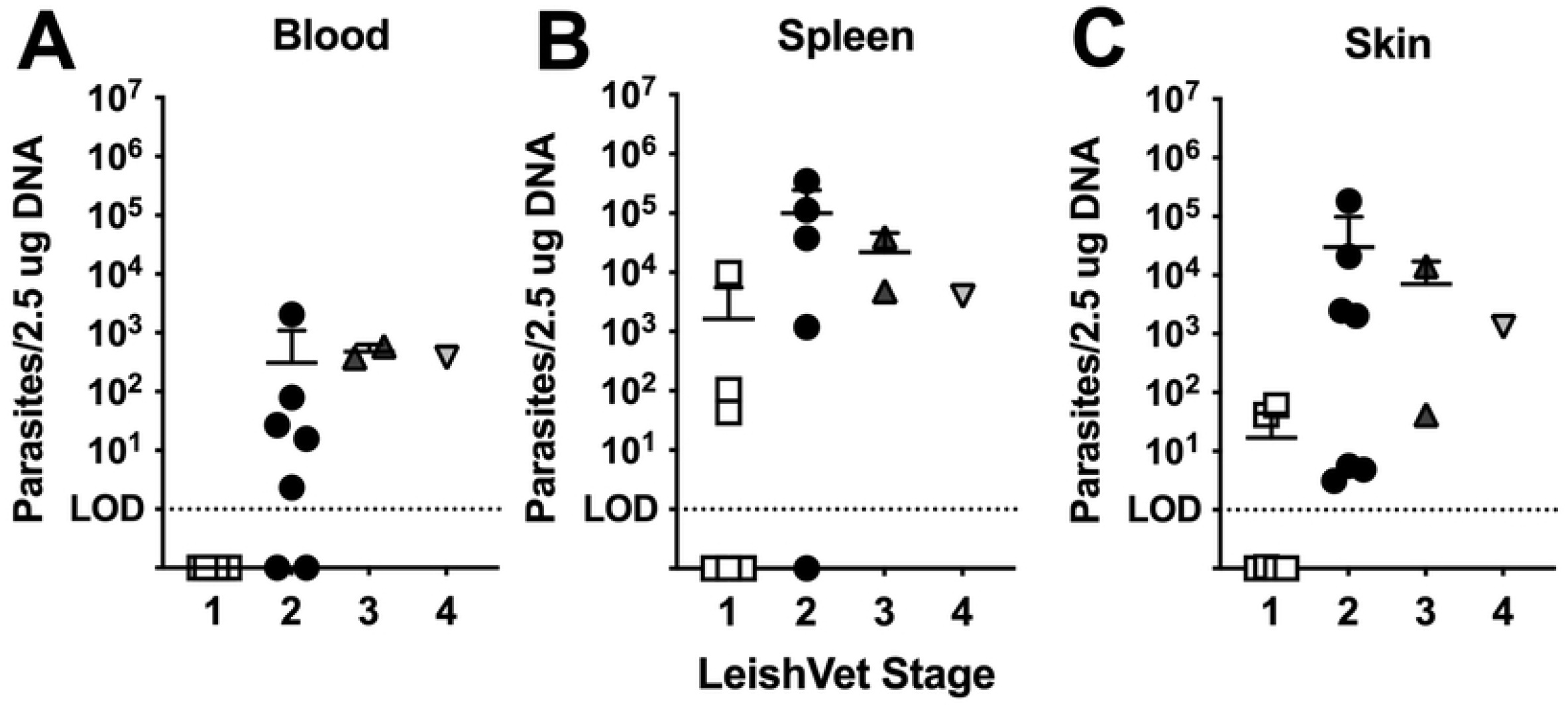
Vertically infected dogs harbor significant skin *Leishmania* burden. Calculated *L. infantum* parasite burden in (A) blood (p=0.004; n=16), (B) spleen (p=0.153; n=14), or (C) skin (p=0.072; n=16) from dogs at indicated LeishVet clinical stage of disease. Each symbol represents one dog. For skin, the average burden of all sampled skin sites per dog is shown. P-values calculated via Kruskal-Wallis ANOVA.

Real Time-qPCR for *Leishmania* DNA in peripheral blood samples revealed none of the LeishVet stage 1 dogs had detectable parasitemia, while 71.4% of LeishVet stage 2 dogs, and 100% of LeishVet stage 3 and 4 dogs were parasitemic, with an average blood burden of 312, 480, and 380 parasite equivalents/2.5 ug DNA, respectively (Figure 1A).

The spleen is a major target organ of *Leishmania* infection. When analyzing splenic parasite load we observed 50% of LeishVet stage 1, 75% of LeishVet stage 2, and 100% of LeishVet stage 3 and stage 4 dogs had detectable parasitism with average burdens of 1.64×10^3^, 9.98×10^4^, 2.2×10^4^, and 4×10^3^ parasite equivalents/2.5 ug DNA, respectively (Figure 1B). As expected, the quantified parasite load in the spleen was higher than observed in the peripheral blood.

To investigate skin parasite burden, we assayed between two and four unique skin biopsy sites per dog. The parasite quantification shown in Figure 1C is the average burden across all biopsy sites/dog. Individual skin biopsy burden and parasite burden difference by biopsy location are shown in Supplemental Figure S1. In vertically infected dogs, we observed 50% of LeishVet stage 1 and 100% of LeishVet stage 2, stage 3, and stage 4 dogs had detectable skin infection, with average skin burdens of 17.1, 3×10^4^, 7.1×10^3^, and 1.3×10^3^ parasite equivalents/2.5 ug DNA, respectively (Figure 1C). Despite never receiving intradermal parasites from a vector, the skin of vertically infected dogs contained a substantial parasite burden.

### Skin parasite burden correlates with systemic parasite load

Parasitemia has long been considered a focal determinant of *Leishmania* vector transmission potential, therefore we were interested in establishing whether skin parasite burden from vertically infected dogs correlated with parasitemia. We found a strong significant positive correlation (p=0.002, r=0.75) between skin parasite burden and corresponding parasitemia (Figure 2A). Despite this, the burden of parasites in the skin was consistently higher than the blood burden detected for every case except one (p=0.0046, Figure 2B). Skin parasite burden also significantly positively correlated with splenic parasite burden (p=0.001, r=0.79, Figure 2C). Moreover, in a paired analysis, skin and splenic parasite load were not significantly different (p=0.18), demonstrating that the skin accumulates parasites to a similar extent as the spleen. This is remarkable, because the spleen is one of the most highly parasitized tissues in VL.

**Figure 2.**
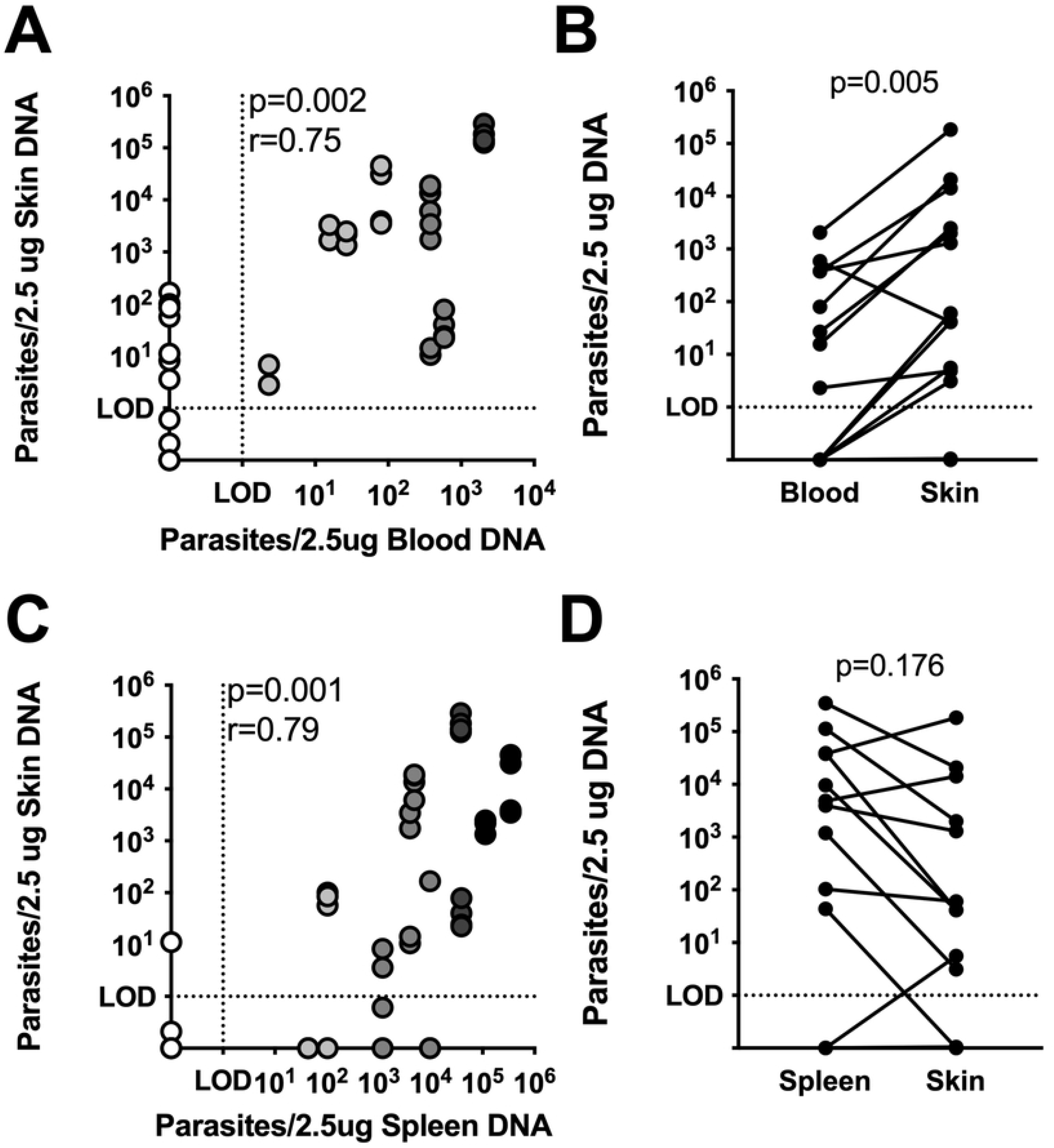
Skin parasite burden correlates with systemic parasite load. Spearman’s correlation (A) and paired Wilcoxon test (B) between skin parasite load and blood parasite load from the same animals (n=16). Spearman’s correlation (C) and paired Wilcoxon test (D) between skin parasite load and splenic parasite load from the same animal (n=14). r, Spearman correlation coefficient.

### Dogs with moderate clinical disease show the highest parasite transmissibility to sand flies

Although it was demonstrated that vertically infected dogs could transmit parasites to sand flies [23], it is poorly understood how host factors predict enhanced transmission from this group. For the first time, xenodiagnosis was performed on 16 vertically infected dogs across progressive stages of VL and infection severity. There was no significant difference in the ability of sand flies to feed on dogs (% female sand flies containing blood meal) based on clinical stage (Figure 3A). After feeding, DNA was isolated from individual blood fed sand flies and analyzed against a standard curve to quantify *Leishmania* (Figure S2A).

**Figure 3.**
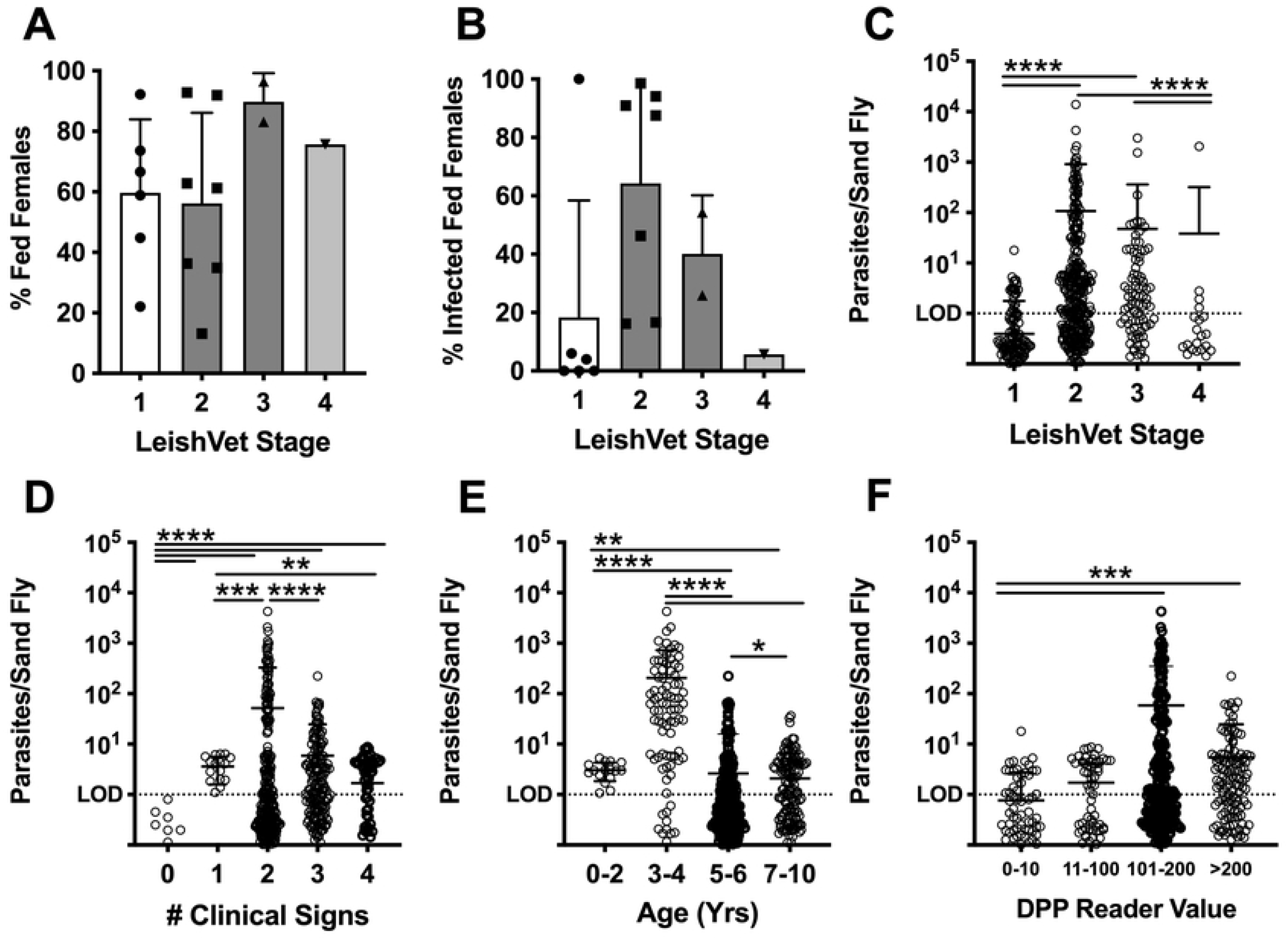
Higher parasite uptake among sand flies fed on dogs with moderate disease. (A) Frequency of sand flies containing a blood meal and (B) frequency of blood meal containing sand flies containing >1 calculated parasite after xenodiagnosis on dogs at indicated LeishVet clinical stages. (C-F) Calculated number of parasites per individual sand fly after xenodiagnosis on dogs with indicated LeishVet clinical stage (C), number of VL clinical signs (D), age range (E), or DPP serological value (F). Kruskal Wallis ANOVA with Dunn’s post-test. **p<0.01; ***p<0.001; ****p<0.0001.

Although flies successfully fed on all dogs at a similar rate, we measured variable parasite equivalents in each sand fly after feeding. (Figure S2B). Average parasite quantification post-feeding was similar between sand flies fed on pinnae or axillary feeding sites (Figure S2C). Interestingly, the highest percentage of infected sand flies (% of blood fed flies containing >1 parasite equivalent) was found after feeding on LeishVet stage 2 dogs, compared to more clinically apparent stage 3 and 4 dogs (Figure 3B, S2B). Similarly, quantification of parasites taken up by individual sand flies showed dogs at LeishVet stage 2 and 3 took up significantly more parasites than LeishVet stage1, subclinical disease, or stage 4, severe disease, dogs (Figure 3C). As some previous studies have observed increased transmission from symptomatic dogs, we also stratified sand fly parasite uptake by number of VL clinical signs. We found dogs with any clinical signs of disease led to more sand fly parasite uptake than was seen by dogs with zero clinical signs of disease. However, similar to the LeishVet stage, parasite uptake did not directly correlate with increasing number of clinical signs (Figure 3D, p=0.24).

The age range of dogs with the highest average parasite uptake per sand fly in this cohort was 3-4 years old (Figure 3E). Serological responses to *L. infantum* rK28 antigen rise with increasing disease severity [18], therefore we compared DPP®CVL reader value correlation with parasite uptake by sand flies and found seronegative dogs (Reader Value 0-10) had the lowest transmission to sand flies, while dogs in the two highest serologically positive bins resulted in significantly higher transmission to sand flies (Figure 3F).

Therefore, none of these factors (clinical severity, age, or serology) directly correlated with parasite transmission to sand flies. Instead, we observed a “Goldilocks” effect, where dogs with moderate disease resulted in the highest rate of sand fly infection and highest parasite burden in infected sand flies.

### Skin parasite burden is the best correlate of successful parasite transmission to sand flies

Parasitemia is known to correlate with *Leishmania* parasite transmission to sand flies. However, in almost every subject, the skin of vertically infected hounds contained a higher parasite burden compared to matched blood samples (Figure 2B). Therefore, we were very interested to know if skin parasite load in this cohort would correlate with transmission to sand flies. In Figure 4A, there was a strong positive correlation (Spearman r=0.68, p=0.005) between skin parasite burden and sand fly parasite uptake. Parasite load in the blood (Spearman r=0.64, p=0.009) and spleen (Spearman r=0.52, p=0.058) also positively correlated with sand fly parasite uptake. However, the parasite burden in the skin of our cohort showed both the strongest correlation coefficient and the most significant association with parasite transmission to sand flies by xenodiagnosis.

**Figure 4.**
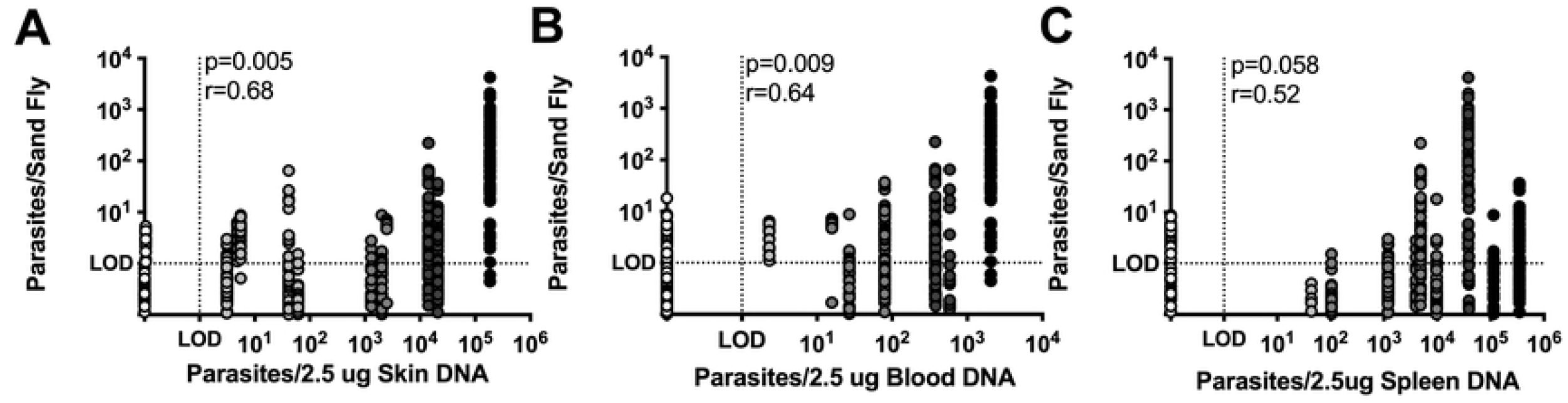
Skin parasite burden is best correlate of parasite transmission to sand flies. Correlation between calculated parasite uptake by individual sand flies after xenodiagnosis and paired (A) average skin parasite burden (n=16), (B) blood parasite burden (n=16), (C) or splenic parasite burden (n=14). Spearman correlation p-value and correlation coefficient (r) are shown.

## DISCUSSION

This study is the first investigation into parasite dermotropism and its relationship to xenodiagnosis in a vertically infected canine VL cohort. Our data supports significant parasite burdens in sand fly accessible tissues, such as blood and particularly skin, as the driving factor determining infectiousness of *L. infantum*-infected dogs. Skin parasite burden was more important than the dog’s clinical status. We found *L. infantum* accumulated in vertically infected dog skin that were significantly higher than those measured in the blood and approaching parasite burdens found in splenic tissue.

Dermal parasite load also differentiates human hosts contributing to the infectious sand fly pool. In people with post-Kala Azar dermal leishmaniasis (PKDL) due to *L. donovani*, which causes VL in India and east Africa, skin parasite load predicted infectiousness to *Phlebotomus argentipes* sand flies by xenodiagnosis [24, 25]. PKDL patients had relatively high skin parasite loads, but blood parasite burdens below the threshold of detection and were still infectious to sand flies via xenodiagnosis [25]. In this study, seropositive but subclinical individuals were not infectious to the vector, similar to our findings of subclinical LeishVet stage 1 dogs. This indicates that it is skin parasite burden, more so than parasitemia, that drives transmissibility of *Leishmania donovani* complex parasites from a mammalian host.

The mechanisms *Leishmania* use to traffic to and seed dermal tissues have not been fully elucidated. It was demonstrated that intravenously administered *Leishmania* parasites trafficked to the skin of B6.*Rag2*^-/-^mice, establishing macro- and microscopic pockets of skin resident infection [13]. This ‘patchy’ parasite distribution better predicted the variable parasite uptake observed in sand flies after xenodiagnosis compared to mathematical models assuming a homogenous parasite source from blood [13]. Our study biopsied multiple skin sites and found limited variability in skin parasite burden from the same subject at the macro level (Figure S1). Thus the microscopic parasite burden at the sand fly bite site seems to influence transmission.

The proficient *Leishmania* dermotropism we found in this population of dogs with vertically transmissible parasites may be evolutionarily advantageous to facilitate vector transmission. Changes in the volatile organic compound profile, or odorous compounds potentially detectable by sand flies, in the hair of dogs from an *L. infantum* endemic area had high sensitivity and specificity for identifying infected dogs [26, 27]. Indeed, an olfactometer bioassay found female *Lu. longipalpis* sand flies were significantly more attracted to hamsters with established *L. infantum* infections [28]. Therefore, a high level of parasites in skin may lead to biochemical changes attractive to the vector, enhancing feeding efficiency and transmission.

Our data agrees with other findings that xenodiagnosis is not driven by symptomatic status. However, we used a higher resolution scale to stage disease, which revealed an interesting dynamic. Moderately diseased dogs (LeishVet levels 2-3) were the most infectious to *Lu. longipalpis* sand flies compared to dogs with mild (LeishVet level 1) or severe disease (LeishVet level 4). This may explain conflicting reports of xenodiagnosis association with clinical signs [7, 8, 10-12, 29, 30]. Dogs with moderate disease may fall into different asymptomatic or symptomatic categories depending on the classification system, and this highlights the need for more standardized staging criteria.

It is interesting to consider why parasites from dogs with severe VL disease may not be as transmissible. Here, *Lu. longipalpis* sand flies successfully fed on a LeishVet stage 4 dog but took up relatively few parasites. LeishVet level 3 and 4 dogs were defined by compromised renal function, which can lead to accumulation of nitrogenous blood products such as blood urea nitrogen as well as elevated breakdown products like creatinine (Table S2) [19]. These waste products may interfere with blood cell integrity, and parasite viability within the sand fly midgut. The sand flies fed on a LeishVet stage 4 dog in this study were observed containing relatively dark, precipitated blood meals after 48 hrs of incubation, which may indicate altered blood meal digestion. However, the sample size of LeishVet level 3-4 subjects in this study was low and xenodiagnosis using more dogs in this range is necessary to confirm this relationship.

Diagnosing VL early in the infectious course in dogs is difficult due to the non-specific nature of initial clinical signs and clinicopathological results. Interestingly we observed several dogs with undetectable parasitemia already had detectable parasites in the skin by qPCR. This could implicate skin biopsy as a more sensitive diagnostic in these instances. Importantly, some sand flies fed on these dogs were able to successfully take up parasites, demonstrating that these early-stage dogs are also infectious to sand flies as seen in our experiments, despite having a low blood burden.

Due to institutional requirements, this study was not able to capture whether all infected sand flies would go on to develop high levels of infection and virulent metacyclic promastigotes. Longer incubation times are needed for parasites to undergo logarithmic replication and metacyclogenesis in the sand fly. We expect, with longer incubation times, the correlation between skin parasite load and sand fly infection level would be even stronger. Additionally, Serafim *et al*. showed sequential blood meals significantly boost parasite replication in the sand fly midgut [31]. How skin parasite burden affects sand fly xenotransmission is a topic for future investigation.

Xenodiagnosis is the most accurate way to ascertain reservoir infectivity to vectors, however it is not trivial to perform. More easily measured correlates of xenodiagnosis need to be identified for practical application. Our data supports skin parasite load as a surrogate marker of dogs with *L. infantum* transmission potential from both horizontally and vertically infected dogs. From a public health perspective, this study reinforces and highlights the real contribution vertically infected dogs present to the infectious burden of the domestic reservoir. Within this cohort, vertically infected dogs have a basic reproductive number (R_θ_) of ∼5, which may vary by breed and litter size [4]. Vertical transmission can occur independently of vector-targeted interventions like indoor residual spraying and insecticide collars. Thus, neutering *Leishmania* infected dogs and the offspring of infected dogs regardless of diagnostic status, is critical to prevent vertical transmission and must be incorporated into *Leishmania* control strategies.

## ACKNOWLEDGEMENTS

We would like to thank the participating animal caretakers who donated subjects to the University of Iowa for this work. We also acknowledge the University of Iowa Comparative Pathology Core technicians for their valuable necropsy assistance. This research was partially supported by the Intramural Research Program of the NIH, National Institute of Allergy and Infectious Diseases. This work was completed while Breanna Scorza was supported by the University of Iowa Interdisciplinary Immunology Postdoctoral Training Grant NIH/NIAID T32AI007260.

## SUPPLEMENTAL FIGURE LEGENDS

**Supplemental Table S1. Cohort demographics**.

Overview of the age in years, sex, and LeishVet status of xenodiagnosis cohort.

**Supplemental Table S2. Cohort Bloodwork**.

Overview of serum chemistry and blood count findings for dogs in each LeishVet clinical grouping. Bolded values indicate the mean is outside of the normal reference range.

**Supplemental Figure S1. Skin parasite burden overview**. (A-B) Calculated *L. infantum* parasite burden in skin from dogs at indicated LeishVet clinical stage of disease. (A) Each dot represents one skin biopsy. 2-4 skin biopsies from each dog are shown. Kruskal-Wallis with Dunn’s post-test. (B) Each dot represents one skin biopsy, separated by subject. (C) Calculated *L. infantum* parasite burden in skin from biopsies collected at either the axillary region or pinna. Each dot represents the average of 1-2 skin biopsies taken at the indicated site from a single dog. Paired Wilcoxon test. Mean and standard deviation are shown for all.

**Supplemental Figure S2. Calculation of *Leishmania* uptake by individual Sand Flies post xenodiagnosis**. (A) Standard curve derived from *Leishmania* parasite spiked sand flies Real Time qPCR. (B) Parasite quantification results of all blood-meal containing female sand flies obtained after xenodiagnosis by LeishVet clinical stage. (C) No difference in mean parasite number uptake in blood fed sand flies based on anatomical feeding location via Wilcoxon paired analysis.

